# Novel AI-powered computational method using tensor decomposition for identification of common optimal bin sizes when integrating multiple Hi-C datasets

**DOI:** 10.1101/2024.09.29.615651

**Authors:** Y-H Taguchi, Turki Turki

## Abstract

Identifying the optimal bin sizes (or resolutions) for the integration of multiple Hi-C datasets is a challenge due to the fact that bin sizes must be common over multiple datasets. By contrast, the dependence of quality upon bin sizes can vary from dataset to dataset. Moreover, common structures should not be sought in bin sizes smaller than the optimal bin sizes, below which common structure cannot be the primary structure any more even after increasing the number of mapped short reads per bin. In this case, there are no common structures at finer resolutions, suggesting that individual Hi-C datasets may have to be analyzed separately in the bin sizes smaller than the optimal one. Thus, quality assessments of individual datasets have a limited ability to determine the best bin size for all datasets. In this study, we propose a novel application of tensor decomposition (TD) based unsupervised feature extraction (FE) to choose the optimal bin sizes for the integration of multiple Hi-C datasets. TD-based unsupervised FE exhibit phase transition-like phenomena through which the smallest possible bin size (or the highest resolution) can be automatically estimated empirically, without the need to manually set a threshold value for the integration of multiple Hi-C datasets, retrieved from GEO with GEO ID, GSE260760 and GSE255264. To our knowledge, ours is the first one that can optimize bin sizes over multiple Hi-C profiles without any tunable parameters.

## 1 Introduction

The use of optimal bin sizes for Hi-C datasets is critical due to the significant effects that bin sizes can have on outcomes. Bin sizes that are too large (i.e., low resolution) might lead us to overlook biologically important factors that can be seen only at finer resolutions. On the other hand, excessively small bin sizes (i.e., high resolution) might result in a different problem; if short reads that are not large enough are mapped to the individual bins because of an increase in the number of bins, a small number of short reads mapped to a bin causes fluctuations that might prevent us from observing the biologically important structure. Thus, there is a trade-off between resolution and fluctuation. When suppressing sufficiently small fluctuations, it is difficult to achieve a sufficiently high resolution to capture biologically important fine structures. This highlights the need to estimate just how small bin sizes can be without inducing excessively large fluctuations.

Determining the optimal bin size in Hi-C data analysis is crucial, as it directly influences the resolution and quality of detected chromatin interactions. Recent advancements have introduced methods to enhance bin size selection, thereby improving data interpretation. One notable approach is deDoc [1], introduced by Li et al. in 2018. This method selects the bin size at which the structural entropy of the Hi-C matrix reaches a stable minimum, aiming to decode topologically associating domains (TADs) even with ultra-low resolution data. Another significant contribution is QuASAR (Quality Assessment of Spatial Arrangement Reproducibility) [2], developed by Sauria and Taylor. QuASAR assesses the quality of Hi-C data by comparing replicate scores to determine the maximum usable resolution, ensuring that the chosen bin size reflects reproducible spatial arrangements. Additionally, HiCPlus [3], developed by Zhang et al. in 2018, employs deep convolutional neural networks to enhance Hi-C data resolution. Remarkably, it can impute high-resolution Hi-C matrices using only a fraction (1/16) of the original sequencing reads, effectively allowing for smaller bin sizes without compromising data quality. These methodologies represent significant strides in optimizing bin size selection for Hi-C data, facilitating more accurate and efficient analysis of chromatin architecture.

The difficulty in estimating optimal bin sizes increases when integrating multiple Hi-C datasets since an optimal bin size needs to be found for all datasets, in addition to the need to identify an optimal bin size for the individual Hi-C datasets. In reality, it is difficult to know whether the strategy by which we can determine the smallest bin sizes under a reasonable amount of fluctuation for multiple Hi-C datasets simultaneously when integrating them is successful. For example, although Yardımcı et al. [4] performed quality assessment of various tools, HiCRep [5], GenomeDISCO [6], HiC-Spector [7], and QuASAR-Rep [2] to process multiple Hi-C datasets, their conclusions were “we do not recommend using reproducibility scores to attempt to select an appropriate resolution.” The one tool that has been historically used to process multiple Hi-C datasets, namely HSA [8], is no longer available. And although multiHiCcompare [9] deals with multiple Hi-C datasets, it can only be used to perform pairwise comparisons. Thus, it cannot be used to simultaneously determine the optimal bin sizes for multiple datasets. Similarly, although Dedoc2 [10] can be used to determine the optimal bin size using entropy, it can only be applied to a single Hi-C dataset (i.e., to individual single cells) and cannot be applied in an integrated manner. As these examples show, although tools exist that can be used to find the optimal bin sizes for multiple Hi-C datasets, to the best of our knowledge, none of these enable determining the optimal bin sizes over multiple Hi-C datasets when these are integrated. To resolve this problem, we propose a novel application of TD-based unsupervised FE [11], which allows us to observe a phase transition through which the optimal bin sizes can be determined without specifying a threshold value (Fig. 1). To the best of our knowledge, this represents the first successful attempt to determine optimal bin sizes for multiple Hi-C datasets simultaneously when integrating them.

**Fig. 1.**
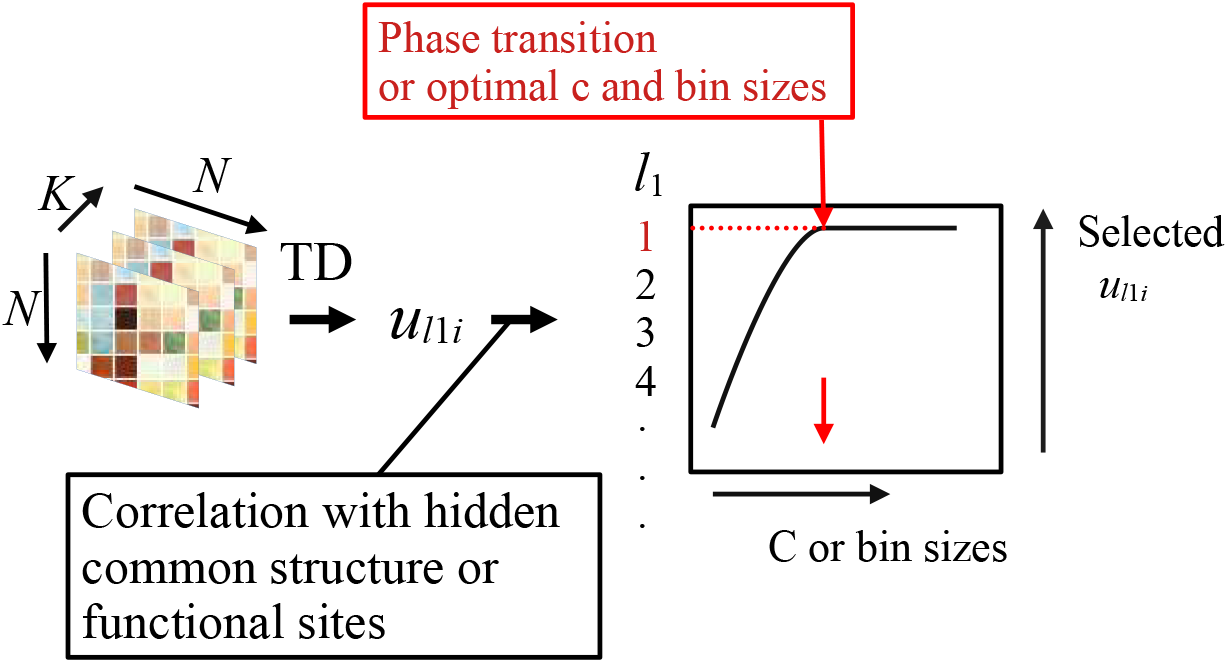
Starting from *K* Hi-C datasets or synthetic data represented as *N* × *N* matrix, a tensor *x*_*ii*′*k*_ ∈ ℝ^*N ×N ×K*^ is generated. Applying TD to *x*_*ii*′*k*_ gives 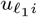, which is attributed to *i*th bins or features. As long as *c*, which is a contribution of a hidden common structure, or the bin sizes are large enough, *ℓ*_1_ = 1 is correlated with a hidden common structure or functional sites. Optimal *c* or bin size is decided at the phase transition after which *ℓ*_1_ = 1 is no longer correlated with a hidden common structure or functional sites.

There are multiple reasons why TD-based unsupervised FE is suitable to address this problem, i.e., optimal bin sizes for multiple Hi-C data sets. In principal, TD-based unsupervised FE was applicable to wide ranged genomic problems [11], including gene expression, histone modification, DNA methylation, chromtine structures, and so on. Thus, we can expect that TD-based unsupervised FE can deal with Hi-C profiles as well. Especially, since we recently successfully have applied TD-based unsupervised FE to protein-protein interaction (PPI) [12–14], there are no reasons that we do not apply TD-based unsupervised FE to Hi-C profiles, since Hi-C and PPI are interaction between genomic loci in some sense.

## 2 Results

### 2.1 Synthetic data

First, we investigated the singular value vector, 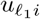, attributed to the *i*th feature, which corresponds to the *i*th bin when we consider real Hi-C data. This is associated with a hidden common (clustering) structure, *y*_*i*_ (see Methods) embedded into the synthetic dataset where there are 1,000 features (*i*), the top 10 of which are clustered with each other (for more details, see Methods). Figure 2 shows the dependence of the selected *ℓ*_1_ on *c*, which represents the contribution of the hidden common structure among all datasets (see Methods). Although *ℓ*_1_ = 1, which has the largest contribution, is always most highly correlated with a common structure for a larger *c*, the frequency of selecting *ℓ*_1_ other than 1 increases as *c* decreases. Additionally, although *ℓ*_3_, which is most highly associated with the selected *ℓ*_1_, is always 1, which corresponds to simple averaging over the samples (Fig. 3), for large enough *c*, other *ℓ*_3_s than 1 will be selected as *c* decreases (Fig. 4). In reality, although the correlation between the selected 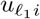 and hidden common structure *y*_*i*_ remains equal to one for a larger *c*, it decrease as *c* decreases below a certain threshold value (Fig. 5); for smaller values of *c*, the correlations are no longer significant (Fig. 6).

**Fig. 2.**
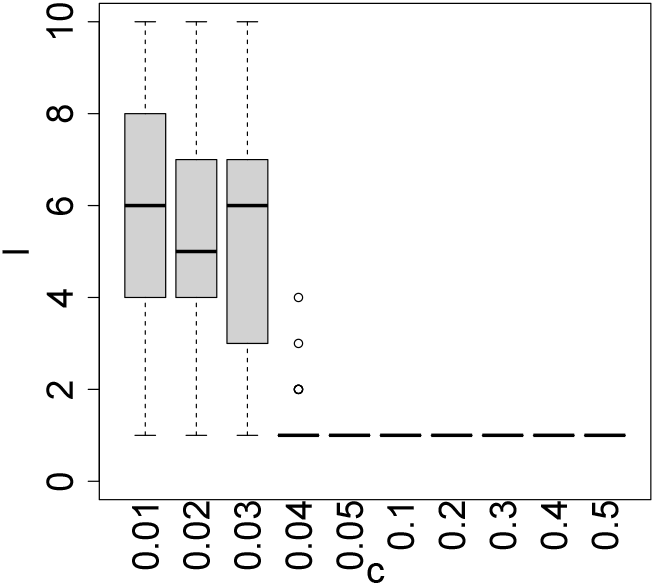
The boxplot of *ℓ*_1_ selected upon *c*

**Fig. 3.**
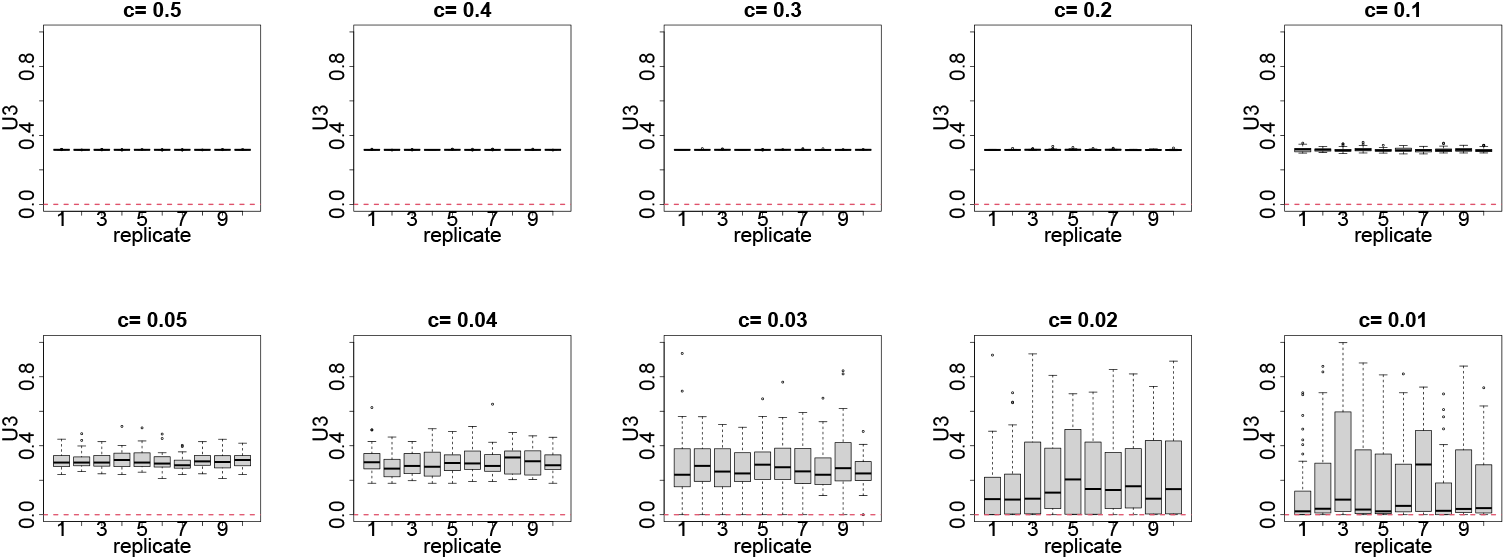
Boxplot of 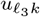 over replicates (1 ≤ *k* ≤ 10) for various *c* values. The red horizontal broken line represents 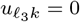

**Fig. 4.**
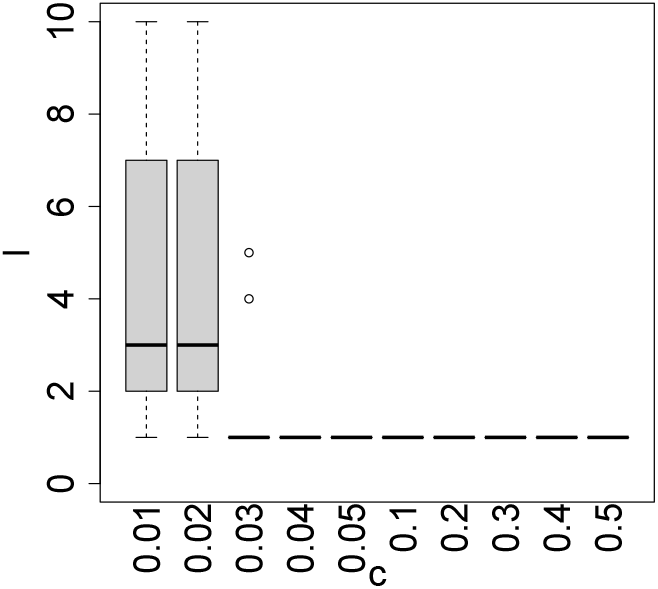
The boxplot of *ℓ*_3_ selected upon *c*

**Fig. 5.**
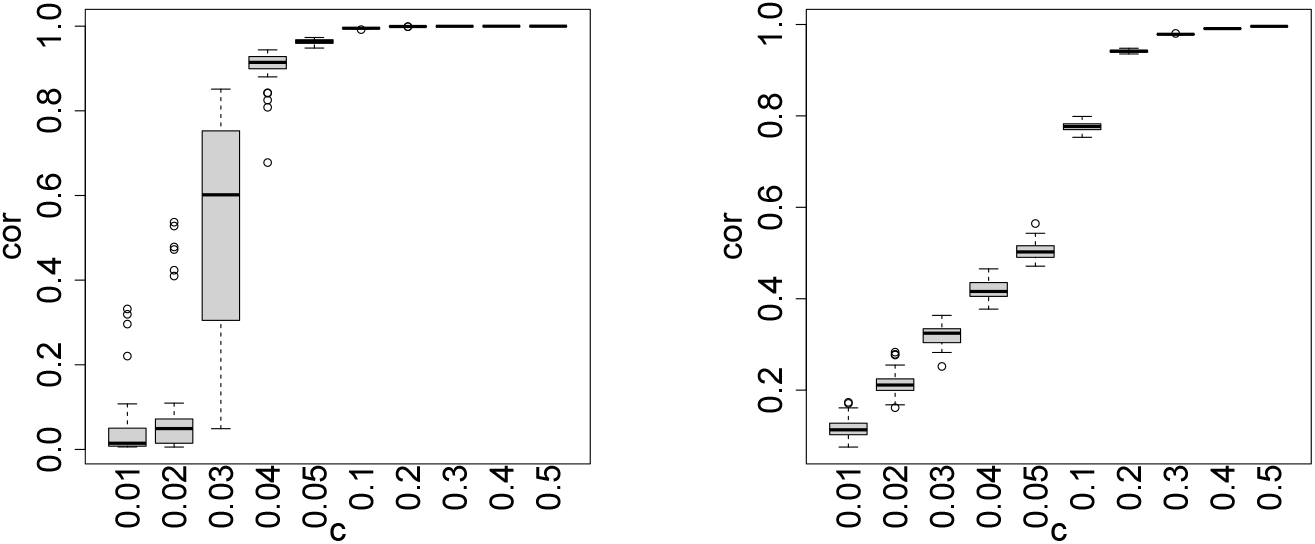
The boxplot of the Pearson’s correlation coefficients between 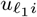 selected (left) or simple average ⟨*x*_*ii*′*k*_ ⟩_*i*_ (right) and a hidden common structure, *y*_*i*_, as a function of *c*

**Fig. 6.**
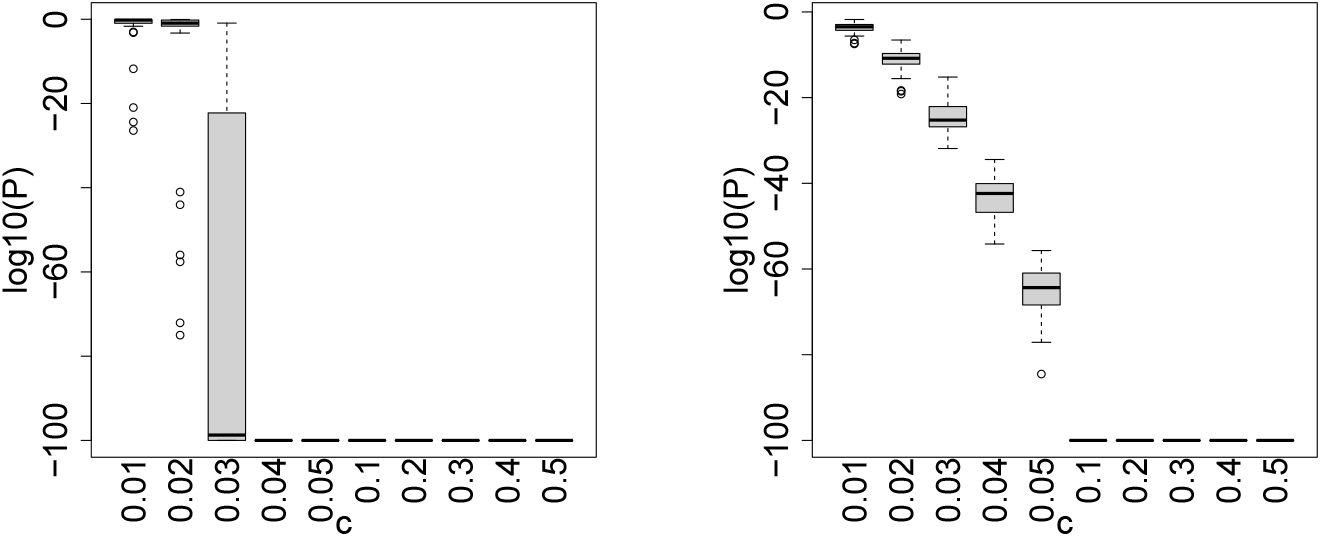
The boxplot of *P* -values in the logarithmic scale associated with the Pearson’s correlation coefficient between 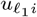 selected (left) or simple average ⟨*x*_*ii*′*k*_⟩_*i*_ (right) and a hidden common structure, *y*_*i*_, as a function of *c. P <* 10^−100^ are truncated to be *P* = 10^−100^.

Lastly, we checked whether the TD-based unsupervised FE correctly selected the 10 features (*i* ≤10) that were interconnected with each other (Table 1). Although FPs (i.e., *i >* 10 but wrongly associated with adjusted *P* -values less than 0.01) heavily fluctuated, TPs (i.e., *i* ≤10 and correctly associated with adjusted *P* -values less than 0.01) monotonically decreased, and FNs (i.e., *i* ≤10 but wrongly associated with adjusted *P* -values larger than 0.01) monotonically increased as *c* decreased after *c* ≤0.05. This is consistent with the observation that a TD-based unsupervised FE cannot capture the hidden common structure *y*_*i*_ if *c* is too small. One could ask why we need TD if a simple average over replicates can achieve a better performance. In response, we can compute a correlation between a simple average ⟨*x*_*ii*′*k*_ ⟩ and a hidden common structure, *y*_*i*_, as well (Fig. 5 and 6) and compare these results with those of TD. From this, it is evident that the TD-based unsupervised FE outperforms the simple average, demonstrating that TD-based unsupervised FE is more worthwhile than the use of a simple average. Based on these observations, the threshold value, below which the hidden common structure is no longer a primary structure, is approximately *c* = 0.05. Although this specific *c* value may not be important, it is important to note that there is a threshold value above which *u*_1*i*_ is always selected and is highly coincident with the hidden common structure. This is analogous to phase transition. It is worth noting that we might be able to use this result for quality assessment; that is, the results obtained by the integration of multiple profiles can be trusted only when they are above the threshold value, and *we do not have to specify the threshold value in advance because the threshold value is automatically decided when u*_1*i*_ *is not correlated with the hidden common structure, y*_*i*_. Nevertheless, in real applications, we cannot know what the hidden common structure is. Thus, due to the fact that we need a hidden common structure in order to compute correlation coefficients, it may not be possible to estimate threshold values. In the following, we demonstrate how to estimate an accurate threshold value using the results obtained by integrating multiple profiles without knowing hidden common structures.

**Table 1.**
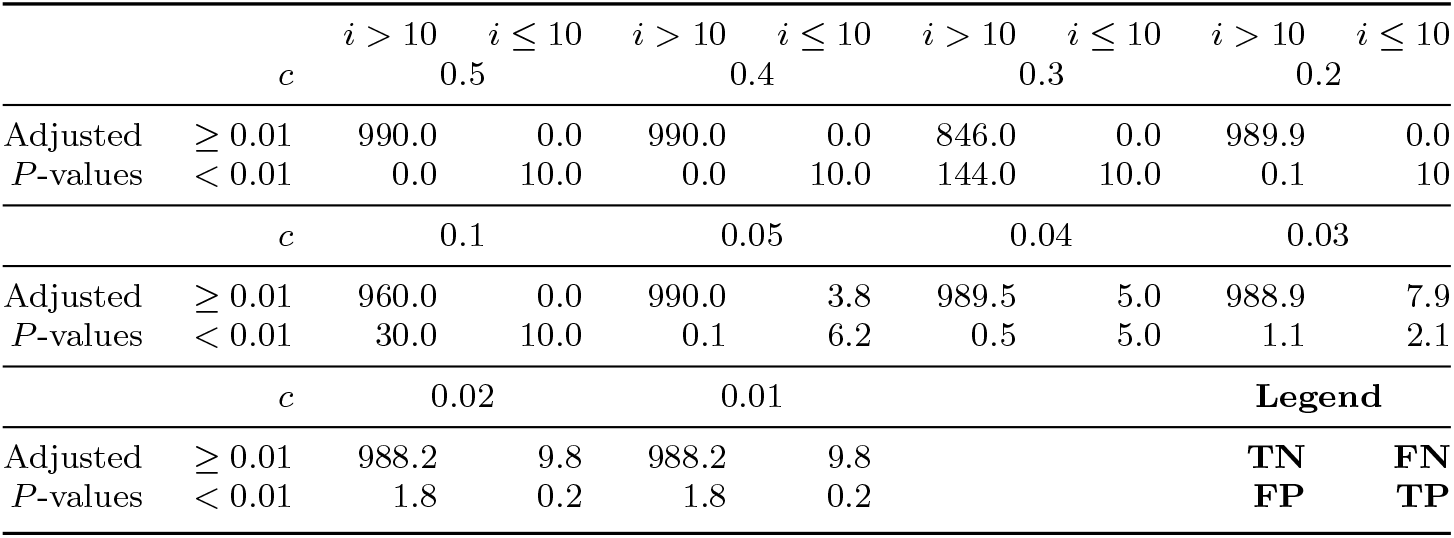
Confusion matrix between sites selected (associated with adjusted *P* -values less than 0.01) by TD-based unsupervised FE and those interconnected sites (*y*_*i*_ = 1, *i* ≤10). The correspondences to TP (true positive), FP (false positive), TN (true negative), and FN (false negatives) are also shown at the end of table.

### 2.2 Hi-C dataset

In this section, the application of TD-based unsupervised FE to real Hi-C datasets and the use of the resulting phase transition for quality assessment, as mentioned above, is described for the selection of optimal bin sizes.

Herein, TD-based unsupervised FE was applied to two Hi-C datasets. Because the hidden common structures are unknown, known functional sites can be employed instead of hidden common structures (Tables 2, 3, 4, and 5). Here are explanation of functional sites. CTCF is the transcription factor, CTCF, binding sites. CTCF transcription factor is know to be related to chromatin structures. Thus, the coincidence with CTCF binding site is good measure to evaluate whether Hi-C profiles capture the functional sites. PLS, pELS, and dELS are know to be overlap with binding sites of TAZ, activation of which is known to result in chromatin architecture alterations [15]. Thus they are also good measutres to evaluate whether Hi-C profiles capture the functional sites.

**Table 2.** Absolute correlations between 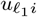 or simple average ⟨*x*_*ii*′*k*_⟩_*i*_ and CTCF binding sites. Bold is used to denote the highest absolute correlation coefficients.

| CTCF<br>PC | 1 | 2 | 5 | $\langle x_{ii'k} \rangle_i$ |
| --- | --- | --- | --- | --- |
| GSE260760 |  |  |  |  |
| **1000000** |  |  |  |  |
| Pearson | **$4.98 \times 10^{-1}$** | $1.23 \times 10^{-1}$ | | $3.36 \times 10^{-1}$ |
| P-value | $1.43 \times 10^{-175}$ | $6.49 \times 10^{-11}$ | | $5.64 \times 10^{-75}$ |
| Spearman | **$5.72 \times 10^{-1}$** | $4.99 \times 10^{-1}$ | | $5.18 \times 10^{-1}$ |
| P-value | $8.34 \times 10^{-244}$ | $7.69 \times 10^{-177}$ | | $8.30 \times 10^{-193}$ |
| **150000** |  |  |  |  |
| Pearson | $5.65 \times 10^{-3}$ | $2.75 \times 10^{-2}$ | **$4.42 \times 10^{-1}$** | $3.14 \times 10^{-1}$ |
| P-value | $4.41 \times 10^{-1}$ | $1.76 \times 10^{-4}$ | 0 | 0 |
| Spearman | $3.66 \times 10^{-1}$ | $5.52 \times 10^{-1}$ | **$5.73 \times 10^{-1}$** | $5.14 \times 10^{-1}$ |
| P-value | 0 | 0 | 0 | 0 |
| GSE255264 |  |  |  |  |
| **1000000** |  |  |  |  |
| Pearson | **$5.09 \times 10^{-1}$** | $3.62 \times 10^{-2}$ | | $4.45 \times 10^{-1}$ |
| P-value | $1.24 \times 10^{-184}$ | $5.52 \times 10^{-2}$ | | $1.62 \times 10^{-136}$ |
| Spearman | **$6.87 \times 10^{-1}$** | $3.15 \times 10^{-1}$ | | $6.35 \times 10^{-1}$ |
| P-value | 0 | $1.10 \times 10^{-65}$ | | 0 |
| **150000** |  |  |  |  |
| Pearson | **$4.46 \times 10^{-1}$** | $4.01 \times 10^{-2}$ | | $3.88 \times 10^{-1}$ |
| P-value | 0 | $4.38 \times 10^{-8}$ | | 0 |
| Spearman | **$6.25 \times 10^{-1}$** | $2.54 \times 10^{-1}$ | | $5.79 \times 10^{-1}$ |
| P-value | 0 | $3.40 \times 10^{-271}$ | | 0 |
| **40000** |  |  |  |  |
| Pearson | $2.88 \times 10^{-3}$ | **$3.89 \times 10^{-1}$** | | $3.32 \times 10^{-1}$ |
| P-value | $4.47 \times 10^{-1}$ | 0 | | 0 |
| Spearman | $4.42 \times 10^{-1}$ | **$5.55 \times 10^{-1}$** | | $5.11 \times 10^{-1}$ |
| P-value | 0 | 0 |  | 0 |

**Table 3.** Absolute correlations between 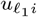 or simple average ⟨*x*_*ii*′*k*_⟩_*i*_ and promoter-like signatures (PLS). Bold is used to denote the highest absolute correlation coefficients.

| PLS<br>PC | 1 | 2 | 5 | $\langle x_{ii'k} \rangle_i$ |
| --- | --- | --- | --- | --- |
| GSE260760 |  |  |  |  |
| **1000000** |  |  |  |  |
| Pearson | **$4.51 \times 10^{-1}$** | $7.08 \times 10^{-2}$ | | $3.25 \times 10^{-1}$ |
| P-value | $1.06 \times 10^{-140}$ | $1.75 \times 10^{-4}$ | | $4.59 \times 10^{-70}$ |
| Spearman | **$6.33 \times 10^{-1}$** | $4.86 \times 10^{-1}$ | | $6.00 \times 10^{-1}$ |
| P-value | 0 | $3.88 \times 10^{-166}$ | | $6.31 \times 10^{-274}$ |
| **150000** |  |  |  |  |
| Pearson | $4.41 \times 10^{-3}$ | $7.91 \times 10^{-3}$ | **$3.32 \times 10^{-1}$** | $2.72 \times 10^{-1}$ |
| P-value | $5.47 \times 10^{-1}$ | $2.81 \times 10^{-1}$ | 0 | 0 |
| Spearman | $2.74 \times 10^{-1}$ | $4.74 \times 10^{-1}$ | **$5.19 \times 10^{-1}$** | $4.77 \times 10^{-1}$ |
| P-value | 0 | 0 | 0 | 0 |
| GSE255264 |  |  |  |  |
| **1000000** |  |  |  |  |
| Pearson | **$6.39 \times 10^{-1}$** | $1.36 \times 10^{-3}$ | | $5.19 \times 10^{-1}$ |
| P-value | 0 | $9.43 \times 10^{-1}$ | | $2.43 \times 10^{-193}$ |
| Spearman | **$7.64 \times 10^{-1}$** | $4.50 \times 10^{-1}$ | | $7.45 \times 10^{-1}$ |
| P-value | 0 | $5.65 \times 10^{-140}$ | | 0 |
| **150000** |  |  |  |  |
| Pearson | **$5.15 \times 10^{-1}$** | $1.50 \times 10^{-3}$ | | $4.17 \times 10^{-1}$ |
| P-value | 0 | $8.38 \times 10^{-1}$ | | 0 |
| Spearman | **$5.87 \times 10^{-1}$** | $2.90 \times 10^{-1}$ | | $5.70 \times 10^{-1}$ |
| P-value | 0 | 0 |  | 0 |
| **40000** |  |  |  |  |
| Pearson | $6.69 \times 10^{-4}$ | **$3.60 \times 10^{-1}$** | | $2.87 \times 10^{-1}$ |
| P-value | $8.60 \times 10^{-1}$ | 0 | | 0 |
| Spearman | $2.75 \times 10^{-1}$ | **$3.93 \times 10^{-1}$** | | $3.68 \times 10^{-1}$ |
| P-value | 0 | 0 |  | 0 |

**Table 4.** Absolute correlations between 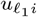 or simple average ⟨*x*_*ii*′*k*_⟩_*i*_ and proximal with enhancer-like signatures (pELS). Bold is used to denote the highest absolute correlation coefficients.

| pELS<br>PC | 1 | 2 | 5 | $\langle x_{ii'k} \rangle_i$ |
| --- | --- | --- | --- | --- |
| GSE260760 |  |  |  |  |
| **1000000** |  |  |  |  |
| Pearson | **$5.43 \times 10^{-1}$** | $8.50 \times 10^{-2}$ | | $3.76 \times 10^{-1}$ |
| P-value | $2.89 \times 10^{-215}$ | $6.47 \times 10^{-6}$ | | $4.77 \times 10^{-95}$ |
| Spearman | **$7.08 \times 10^{-1}$** | $5.46 \times 10^{-1}$ | | $6.75 \times 10^{-1}$ |
| P-value | 0 | $2.00 \times 10^{-217}$ | | 0 |
| **150000** |  |  |  |  |
| Pearson | $4.63 \times 10^{-4}$ | $1.84 \times 10^{-2}$ | **$4.08 \times 10^{-1}$** | $3.23 \times 10^{-1}$ |
| P-value | $9.50 \times 10^{-1}$ | $1.20 \times 10^{-2}$ | 0 | 0 |
| Spearman | $3.32 \times 10^{-1}$ | $5.61 \times 10^{-1}$ | **$6.19 \times 10^{-1}$** | $5.78 \times 10^{-1}$ |
| P-value | 0 | 0 | 0 | 0 |
| GSE255264 |  |  |  |  |
| **1000000** |  |  |  |  |
| Pearson | **$7.01 \times 10^{-1}$** | $3.06 \times 10^{-3}$ | | $5.82 \times 10^{-1}$ |
| P-value | 0 | $8.71 \times 10^{-1}$ | | $1.07 \times 10^{-253}$ |
| Spearman | **$8.22 \times 10^{-1}$** | $4.95 \times 10^{-1}$ | | $8.03 \times 10^{-1}$ |
| P-value | 0 | $3.74 \times 10^{-173}$ | | 0 |
| **150000** |  |  |  |  |
| Pearson | **$5.78 \times 10^{-1}$** | $1.65 \times 10^{-3}$ | | $4.79 \times 10^{-1}$ |
| P-value | 0 | $8.22 \times 10^{-1}$ | | 0 |
| Spearman | **$6.70 \times 10^{-1}$** | $3.56 \times 10^{-1}$ | | $6.54 \times 10^{-1}$ |
| P-value | 0 | 0 |  | 0 |
| **40000** |  |  |  |  |
| Pearson | $9.76 \times 10^{-4}$ | **$4.17 \times 10^{-1}$** | | $3.40 \times 10^{-1}$ |
| P-value | $7.97 \times 10^{-1}$ | 0 | | 0 |
| Spearman | $3.35 \times 10^{-1}$ | **$4.74 \times 10^{-1}$** | | $4.48 \times 10^{-1}$ |
| P-value | 0 | 0 |  | 0 |

**Table 5.** Absolute correlations between 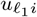 or simple average ⟨*x*_*ii*′*k*_⟩_*i*_ and distal with enhancer-like signatures (dELS). Bold is used to denote the highest absolute correlation coefficients.

| dELS<br>PC | 1 | 2 | 5 | $\langle x_{ii'k} \rangle_i$ |
| --- | --- | --- | --- | --- |
| GSE260760 |  |  |  |  |
| **1000000** |  |  |  |  |
| Pearson | **$8.51 \times 10^{-1}$** | $6.84 \times 10^{-2}$ | | $6.59 \times 10^{-1}$ |
| P-value | 0 | $2.89 \times 10^{-4}$ | | 0 |
| Spearman | **$8.62 \times 10^{-1}$** | $6.45 \times 10^{-1}$ | | $8.44 \times 10^{-1}$ |
| P-value | 0 | 0 |  | 0 |
| **150000** |  |  |  |  |
| Pearson | $6.69 \times 10^{-3}$ | $4.10 \times 10^{-2}$ | **$6.06 \times 10^{-1}$** | $5.69 \times 10^{-1}$ |
| P-value | $3.61 \times 10^{-1}$ | $2.10 \times 10^{-8}$ | 0 | 0 |
| Spearman | $4.54 \times 10^{-1}$ | $7.32 \times 10^{-1}$ | **$8.15 \times 10^{-1}$** | $8.02 \times 10^{-1}$ |
| P-value | 0 | 0 | 0 | 0 |
| GSE255264 |  |  |  |  |
| **1000000** |  |  |  |  |
| Pearson | $7.12 \times 10^{-1}$ | $1.48 \times 10^{-2}$ | | **$7.93 \times 10^{-1}$** |
| P-value | 0 | $4.33 \times 10^{-1}$ | | 0 |
| Spearman | $8.22 \times 10^{-1}$ | $5.88 \times 10^{-1}$ | | **$8.26 \times 10^{-1}$** |
| P-value | 0 | $2.63 \times 10^{-260}$ | | 0 |
| **150000** |  |  |  |  |
| Pearson | $6.05 \times 10^{-1}$ | $9.10 \times 10^{-3}$ | | **$6.63 \times 10^{-1}$** |
| P-value | 0 | $2.14 \times 10^{-1}$ | | 0 |
| Spearman | $7.38 \times 10^{-1}$ | $5.02 \times 10^{-1}$ | | **$7.45 \times 10^{-1}$** |
| P-value | 0 | 0 |  | 0 |
| **40000** |  |  |  |  |
| Pearson | $3.61 \times 10^{-3}$ | $5.11 \times 10^{-1}$ | | **$5.55 \times 10^{-1}$** |
| P-value | $3.40 \times 10^{-1}$ | 0 | | 0 |
| Spearman | $4.80 \times 10^{-1}$ | $6.50 \times 10^{-1}$ | | **$6.51 \times 10^{-1}$** |
| P-value | 0 | 0 |  | 0 |

Since 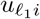 associated with the highest absolute correlation coefficient is almost always more correlated with functional site than the simple average ⟨*x*_*ii*′*k*_⟩_*i*_ excluding dELS for GSE255264, it is reasonable to consider TD-based unsupervised FE as an advanced criteria for the assessment of the quality the Hi-C datasets in order to verify whether the Hi-C data is good enough to capture hidden common structure. A phase transition-like behavior was observed, as expected for the TD-based unsupervised FE applied to a real Hi-C dataset. For GSE260760, *u*_1*i*_ showed the highest absolute correlation with functional sites regardless of the type of functional site for a bin size of 1,000,000, but no longer had a bin size of 150,000. For GSE255264, *u*_1*i*_ showed the highest absolute correlation with functional sites regardless of the type of functional site for bin sizes of 1,000,000 and 150,000, but no longer had a bin size of 40,000. Thus, phase-transition-like phenomena are expected to occur between 1,000,000 and 150,000 for GSE260760 and between 150,000 and 40,000 for GSE255264. These results indicate that bin sizes less than or equal to 150,000 for GSE260760 and those less than or equal to 40,000 for GSE255264 can no longer be regarded as good enough to capture hidden common structures.

### 2.3 Cluster structure detection of Hi-C datasets

Since TD-based unsupervised FE can be used to select bins associated with functional sites, it may also be used to detect common clustering structures within multiple HiC datasets. To evaluate this, we plotted 2424 and 447 *x*_*ii*′*k*_s selected by *u*_1*i*_ and *u*_2*i*_ with bin size of 40,000 for GSE255264, respectively (Fig. 7). Since in this bin size for GSE255264, *u*_1*i*_ is not associated with the highest absolute correlation with functional sites, *u*_1*i*_ is not expected to be associated with the (unknown) hidden common structure. By contrast, *u*_2*i*_ is expected to be associated with the (unknown) hidden common structure since *u*_2*i*_ is associated with the highest absolute correlation with functional sites (Table 2, 3, 4, and 5). As expected, 2424 *x*_*ii*′*k*_s selected by *u*_1*i*_ had a greater sample dependence than 447 *x*_*ii*′*k*_s selected by *u*_2*i*_, since the mean vs SD ratio of 447 *x*_*ii*′*k*_s selected by *u*_2*i*_ was less than that of 2424 *x*_*ii*′*k*_s selected by *u*_1*i*_ (Table 6). This result suggests that for a bin size of 40,000, the (unknown) hidden common structure is no longer a primary part of datasets, and this is in fact a secondary contribution. In reality, although 2424 *x*_*ii*′*k*_s selected by *u*_1*i*_ are widely distributed along the whole genome, the distribution of 447 *x*_*ii*′*k*_s selected by *u*_1*i*_ is restricted to the limited region along the whole genome (Fig. 7). This demonstrates that with a bin size of 40,000, the present dataset for GSE255264 is unable to capture the common structures within the dataset.

**Table 6.** Mean vs SD ratios. Smaller values are indicative of more independent of samples (*k*s). Bold denotes the *ℓ*_1_s most correlated with functional sites

| $u_{\ell_1 i}$ used for selection<br>$\ell_1$ | GSE255264 | | GSE260760 | | |
| --- | --- | --- | --- | --- | --- |
|  | 1 | **2** | 1 | 2 | **5** |
| Mean vs SD ratio | 0.629 | 0.302 | 0.472 | 0.436 | 0.475 |

**Fig. 7.**
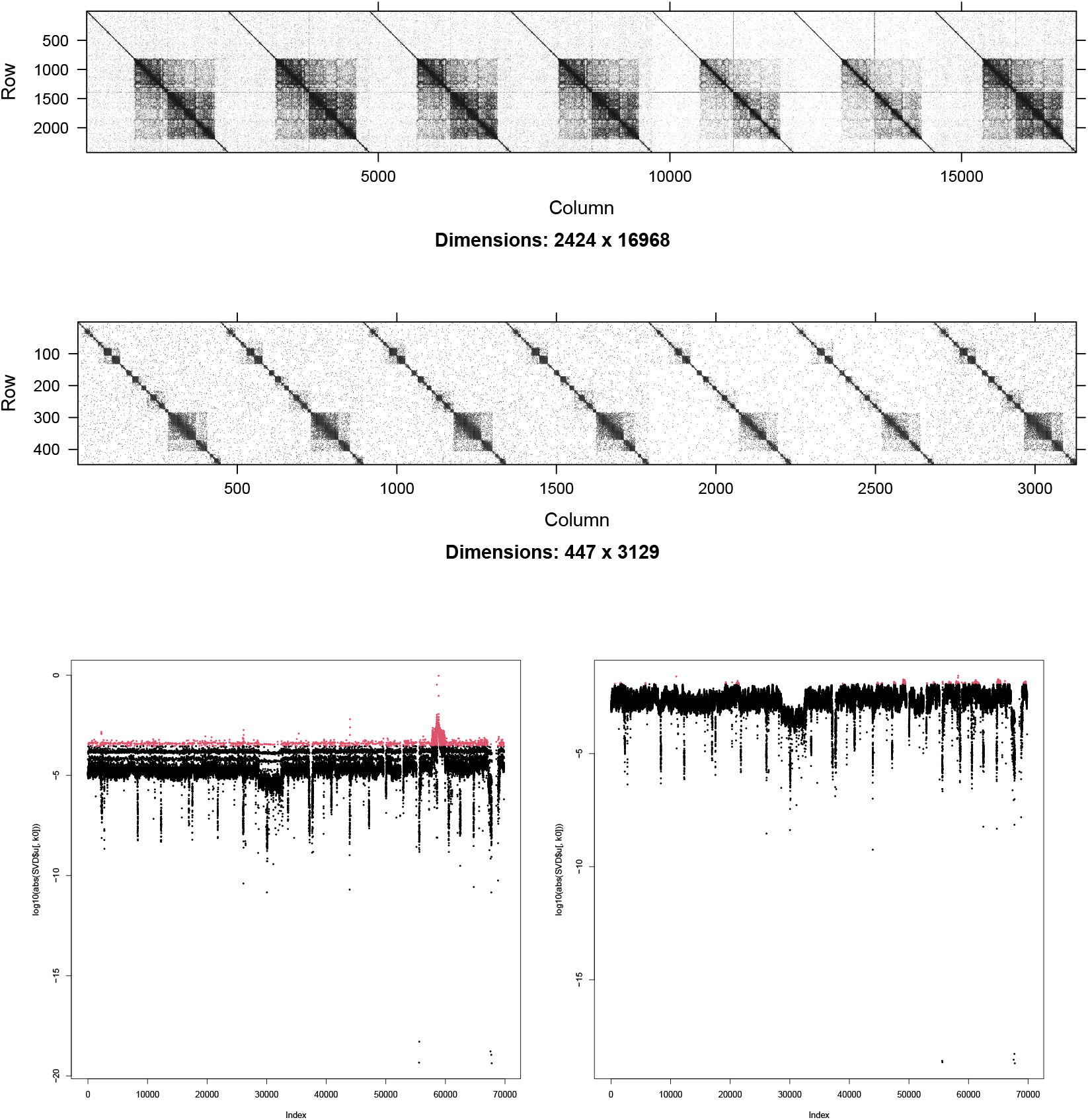
Top: 2424 *x*_*ii*′*k*_s selected by *u*_1*i*_, middle: 447 *x*_*ii*′*k*_s selected by *u*_2*i*_ for GSE255264. From left to right, the seven datasets are aligned horizontally in ascending order of their GSM IDs. Bottom left: log_10_ *u*_1*i*_, bottom right log_10_ *u*_2*i*_. Red indicates the selected ones. The bin sizes are 40,000.

However, for GSE260760 where the bin size of 150,000 is far below the phase transition-like point, even *u*_2*i*_ is associated with the highest absolute correlation with functional sites, *u*_5*i*_ (Table 2, 3, 4, and 5). Fig. 8 shows the 225, 171, and 187 *x*_*ii*′*k*_s selected by *u*_1*i*_, *u*_2*i*_, and *u*_5*i*_. The 187 *x*_*ii*′*k*_s selected by *u*_5*i*_ associated with the highest absolute correlation with functional sites cannot be regarded as more independent of some samples than others, since the Mean vs SD ratios are almost equivalent between *u*_1*i*_, *u*_2*i*_, and *u*_5*i*_ (Table 6). The fact that the number of selected bins was small (Fig. 8, 225, 171 and 187 *x*_*ii*′*k*_s that are less than 1,000) also suggests that the integration was substantially degraded. Based on the above observations, the optimal bin sizes for integrating GSE260760 and GSE255264 were 1,000,000 and 150,000, respectively, within the tested bin sizes. As fewer bin sizes are usually employed to investigate genome structures, we need a greater number of reads and smaller bin sizes as thresholds (i.e., to make phase transition-like phenomena occur at smaller bin sizes). The advantage of this strategy is that we can check whether the number of reads is sufficient before conducting a detailed investigation of the obtained contact map. Alternatively, if we are unable to suppress the appearance of phase transition-like phenomena even after increasing the number of mapped short reads, this suggests that there are no common structures at finer resolutions; in this case, multiple Hi-C datasets will need to be integrated at finer resolutions. The strategy presented here informs this type of decision-making. In conclusion, by replacing the correlation between a hidden common structure and that of functional sites, we obtained the following:

**Fig. 8.**
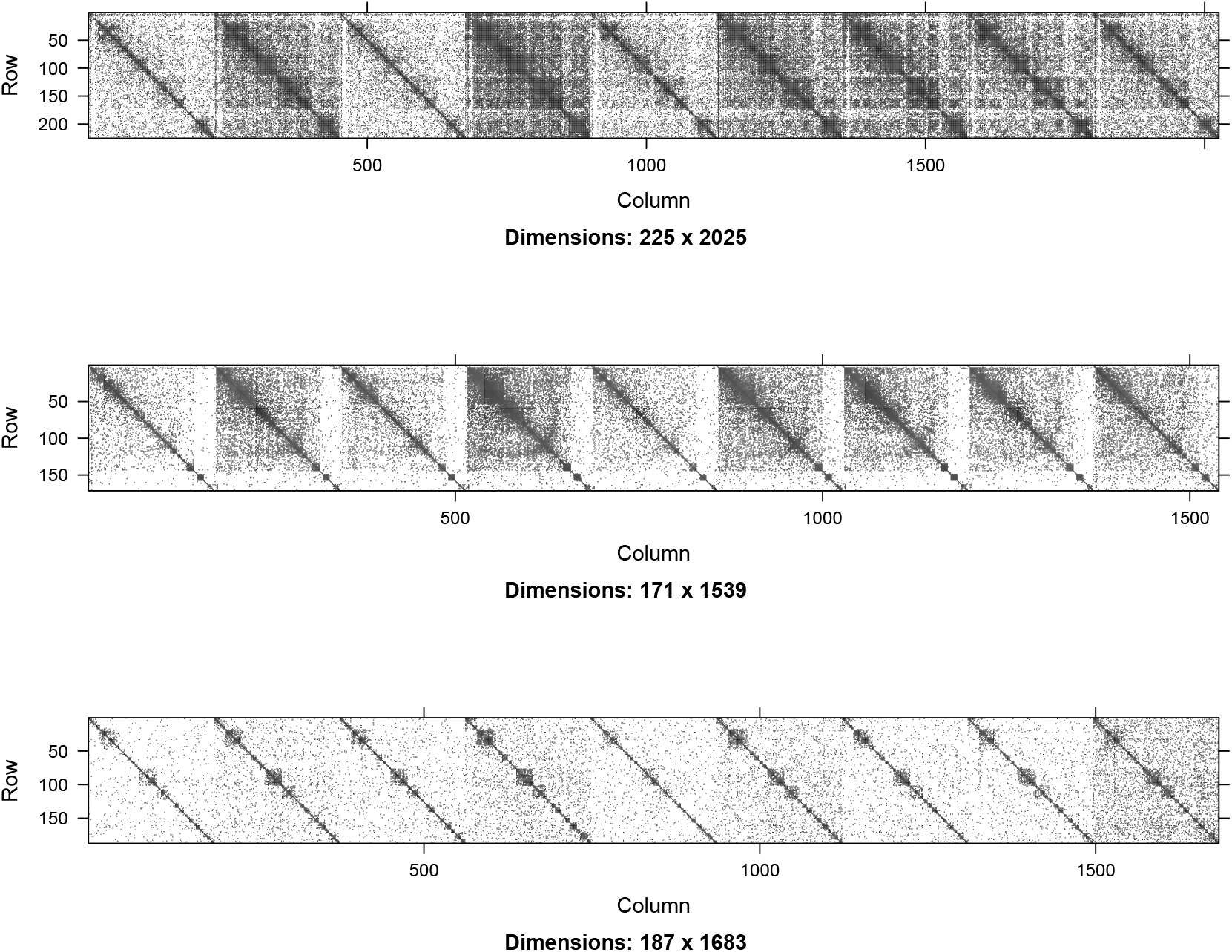
Top: 255 *x*_*ii*′*k*_s selected by *u*_1*i*_; middle: 171 *x*_*ii*′*k*_s selected by *u*_2*i*_; bottom: 187 *x*_*ii*′*k*_s selected by *u*_5*i*_ for GSE260760. The bin sizes are 150,000. From left to right, the nine datasets are aligned horizontally. From left to right, the nine datasets are aligned horizontally in ascending order of the GSM IDs.

1. A realization of the “phase transition-like” phenomena, below which we cannot expect that common structures are primary factors within a dataset (since *u*_1*i*_ is not associated with the highest absolute correlation with functional sites).
2. A more significant correlation with functional sites using TD-based unsupervised FE compared with the simple average ⟨*x*_*ii*′*k*_ ⟩_*i*_.

These results highlight the promising nature of this method.

## 3 Discussion

Changes in bin sizes in the Hi-C data are analogous to changing c, which represents the contribution of the hidden common structure represented by *y*_*i*_ in the synthetic data. As the number of bins is increased and the size of the bin is decreased, the number of reads per individual bin decreases under the condition of a fixed total number of short reads. This results in a small number of reads in individual bins and an inaccurate estimation of the number of reads in individual bins. Thus, as the size of the bins decreases, the fluctuation in the number of reads in the bins increases, resulting in the divergence between the individual datasets. Thus, decreases of bin size should be analogous to those of the parameter “*c*” in the synthetic data. Thus, it is not surprising that the bin sizes decrease, since “*c*” decreases in the synthetic data, as observed through the phase transition-like phenomena. In addition, TD-based unsupervised FE can be used to assess the quality of the Hi-C measurements. Because the number of reads downloaded for individual datasets was fixed at 100,000,000, the number of expected reads in individual bins was also constant. Nevertheless, if the number of interactions differs between datasets, it becomes difficult to capture hidden common structures within multiple Hi-C datasets. For this reason, a decrease in bin size corresponds to a decrease in *c* in the synthetic data. Thus, phase transition-like phenomena can occur, as in the case for synthetic datasets.

As for the comparison with previous studies, to our best knowledge, there are no other previous methods that can derive optimal bin sizes based upon multiple profiles without selecting no threshold values in advance. Although there are some deep-learning based methods [16–19], they are not applicable to Hi-C data as it is. Thus, instead of the performance comparison with other/previous methods, we compare the our results with various criteria generated by one of SOTA, HiCexplolar [20] (Table 7). For the consistency with our analysis, at first, we noticed that bin size 1,000,000, GSE255264, respectively). On the other hand, bin size 150,000 for GSE269769 and 40,000 for GSE255264, which is judged not to be acceptable in our analysis, are associated with sparsity larger than 80 %. Since sparsity less than 80 % is generally supposed to be acceptance condition of Hi-C profile quality, our analysis is primarily coincident with the critetia based upon sparsity. On the other hand, bin side 150,000 for GSE255264, which is judged to be acceptable in our analysis, is associated with sparsity larger than 80 %. Thus, our criteria can have ability to judge the bin size larger than 80% as acceptable. To see what causes the difference between our criterion and that based upon sparsity, we consider other quantity and try to find those distinct between bin size 150,000 for GSE269769 and bin size 150,000 for GSE255264 since the former is judged not to be acceptable in our criteria whereas the latter it judged to be acceptable in our analysis. Then we found that TADs numbers for GSE255264 are significantly larger than those for GSE269769 (*P* -value computed *t* test is 0.03517) and similarity for GSE255264 are significantly less than those for GSE269769 (*P* -value computed *t* test is 7.757× 10^−10^). This is reasonable since our criteria is based upon biological significance. The number of TADs is supposed to be related to other biological features, e.g, CTCF binding. Since our criteria do not require any threshold value to identify optimal bin sizes, for which HiCexplolar requires threshold value of sparsity 80%, our method is superior to the other method. Actually speaking, even if individual profiles are not problematic in the quality, this does not always mean that there are common structures among them. This suggests that we need another criteria apart from quality assessment of for the integrated analysis apart from individual quality assessment. One should also notice that the correlations between ⟨*x*_*ii*′*k*_⟩_*i*_ and functional sited are not distinct at all between bin sizes with sparsity less than 80 % and others. It also suggests that the present strategy is more useful to evaluate optimal bin sizes than simple average.

**Table 7.** Various measures about Hi-C data quality provided by HiCexplorer. CCS:Compartment Clarity Score.

| GSE269769 |  | GSM81238XX |  |  |  |  |  |  |  |
| --- | --- | --- | --- | --- | --- | --- | --- | --- | --- |
| XX | 32 | 33 | 34 | 35 | 36 | 37 | 38 | 39 | 40 |
| 1000000 | (binsize) |  |  |  |  |  |  |  |  |
| Sparsity | 80.03 | 57.04 | 74.32 | 50.48 | 55.89 | 43.67 | 64.58 | 67.18 | 32.91 |
| Coveragy | 3392.37 | 5051.81 | 4102.69 | 5084.6 | 6653.28 | 6347.97 | 4024.74 | 2953.19 | 10791.09 |
| TADs | 151 | 163 | 144 | 170 | 156 | 160 | 160 | 169 | 154 |
| CCS | 0.15 | 0.15 | 0.15 | 0.15 | 0.15 | 0.15 | 0.15 | 0.15 | 0.15 |
| 150000 | (binsize) |  |  |  |  |  |  |  |  |
| Sparsity | 99.18 | 97.68 | 98.97 | 96.97 | 98.21 | 97 | 97.9 | 98.26 | 96.79 |
| Coveragy | 508.8068 | 757.6976 | 615.3432 | 762.6163 | 997.8959 | 952.1036 | 603.6518 | 442.9353 | 1618.507 |
| TADs | 786 | 1220 | 760 | 1282 | 953 | 1190 | 1291 | 1273 | 1062 |
| CCS | 0.06 | 0.06 | 0.05 | 0.06 | 0.06 | 0.06 | 0.06 | 0.06 | 0.06 |
| similarity | 0.715999 | 0.73567 | 0.729842 | 0.747258 | 0.732688 | 0.722467 | 0.727329 | 0.746207 | 0.702418 |
| GSE255264 |  | GSM80677XX |  |  |  |  |  |  |  |
| XX | 73 | 74 | 75 | 77 | 78 | 79 | 80 |  |  |
| 1000000 | (binsize) |  |  |  |  |  |  |  |  |
| Sparsity | 34.16 | 32.43 | 39.87 | 35.21 | 67.53 | 70.44 | 38.8 |  |  |
| Coveragy | 6045.88 | 6861.03 | 6252.25 | 6038.25 | 2759 | 2321.94 | 6217.18 |  |  |
| TADs | 143 | 141 | 150 | 145 | 146 | 144 | 147 |  |  |
| CCS | 0.15 | 0.15 | 0.15 | 0.15 | 0.15 | 0.15 | 0.14 |  |  |
| 150000 | (binsize) |  |  |  |  |  |  |  |  |
| Sparsity | 96.25 | 95.95 | 96.54 | 96.28 | 98.45 | 98.63 | 96.47 |  |  |
| Coveragy | 906.794 | 1029.055 | 937.7463 | 900.4138 | 408.4901 | 348.2573 | 932.4871 |  |  |
| TADs | 1276 | 1277 | 1281 | 1278 | 1258 | 1224 | 1294 |  |  |
| CCS | 0.06 | 0.06 | 0.06 | 0.06 | 0.06 | 0.06 | 0.06 |  |  |
| similarity | 0.695706 | 0.673369 | 0.697345 | 0.685025 | 0.6736 | 0.680826 | 0.669958 |  |  |
| 40000 | (binsize) |  |  |  |  |  |  |  |  |
| Sparsity | 99.63 | 99.59 | 99.64 | 99.62 | 99.84 | 99.86 | 99.64 |  |  |
| Coveragy | 241.8052 | 274.4072 | 250.0589 | 241.5002 | 108.9277 | 92.86609 | 248.6565 |  |  |
| TADs | 3664 | 3843 | 3761 | 3759 | 3355 | 3163 | 3698 |  |  |
| CCS | 0.03 | 0.03 | 0.03 | 0.03 | 0.03 | 0.03 | 0.03 |  |  |

## 4 Conclusion

In this study, TD-based unsupervised FE was applied to multiple Hi-C datasets. We found phase transition-like phenomena that are boundaries if *u*_1*i*_ were those that were most highly correlated with the functional site, as observed in the synthetic data. Thus, we can specify the optimal bin size as the smallest bin size, provided that *u*_1*i*_ is the most highly correlated with the functional site. As a by-product, *u*_1*i*_ is usually more strongly correlated with functional sites than with simple averages over several datasets. Therefore, it can be used as a representative profile. In summary, in this study, a strategy for selecting the optimal bin size was developed when integrating multiple Hi-C datasets. As a by-product, we can have representative profiles, 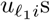,which are more strongly correlated with functional sites than simple averages, ⟨*x*_*ii*′*k*_⟩_*i*_. The weak point of the present methodology is that it is fully unsupervised. Since it does not have any tunable parameters, if it fails, we have to simply give up analysing the data in the present method. Future improvement depends upon the development of TD-based unsupervised FE. For example, Bayesian work frame might be able to improve the method performance.

## 5 Methods

### 5.1 Synthetic data

The synthetic data used in this study was a tensor *x*_*ii*′*k*_ ∈ℝ^1,000*×*1,000*×*10^. Initially, all *x*_*ii*′*k*_s were filled with 0. Then, 100 *x*_*ii*′*k*_s for 1 *≤ i, i*^*′*^ *≤* 10 were filled with 1. In addition to this, 1 was added to *x*_*ii*′*k*_ for 200 randomly selected (*i, i*^*′*^) pairs in individual *k*s (this means that randomly selected pairs are not common between distinct *k*s). The thirty-three *x*_*ii*′*k*_s (replicates) were generated with distinct random seeds. Only 33 replicates were used because only 33 of the 100 replicates were associated with successful HOSVD; the other replicates were eliminated due to singularity.

#### 5.1.1 Hidden common structures

We define *y*_*i*_ as

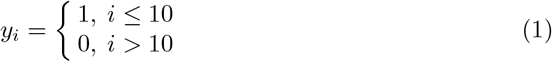

*y*_*i*_ is called a hidden common structure in the analysis of the synthetic data in this study because it is distinct between those interconnected *i*s, regardless of *k*s and the other *i*s.

#### 5.1.2 Correlation with hidden common structures

For synthetic data, the correlations between 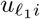 associated with the highest absolute correlation or the simple average

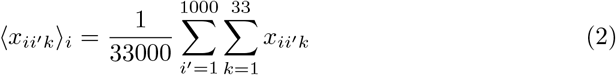

and *y*_*i*_ were computed.

### 5.2 Hi-C datasets

Two Hi-C datasets were analyzed in this study.

#### 5.2.1 GSE260760

Nine Hi-C profiles were downloaded from human knee chondrocytes [21].

#### 5.2.2 GSE255264

Seven AK1 cell line Hi-C profiles [22], that were originally obtained from [23], were downloaded. Although there were nine AK1 datasets, only seven (GSM8067773, GSM8067774, GSM8067775, GSM8067777, GSM8067778, GSM8067779, and GSM8067780) were downloadable.

#### 5.2.3 Downloading fastq files

To have same number of reads for all datasets, “-X 100000000” option, which limits the number of downloaded reads to as many as 108, was added to fastq dump [24].

#### 5.2.4 Mapping reads to the human genome

HiC-Pro [25] was used to map reads to the hg38 human genome, assuming bin sizes of 1,000,000, 500,000, 150,000, 40,000, and 20,000.

#### 5.2.5 Preprocessing

The “raw” matrix files were used for the analyses. *x*_*ii*′*k*_ was normalized as follows

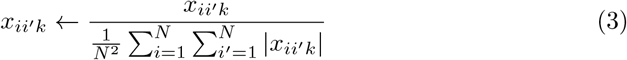

### 5.3 Brief introduction of Tucker decomposition using Higher-order singular value decomposition

There are multiple ways to perform tensor decomposition (TD) [11]. Among them, we employed Tucker decomposition for TD-based unsupervised FE. Tucker decomposition decompose a tensor into a product sum of a small core tensor and matrices whose number is as many as number of indices; this means individual matrices represent bundle of various vectors attributed to each index and dependence upon indices. Usually, there can be several kind of dependence, each column of matrix corresponds to dependence upon index. Usually, TD is expected to reduce the degrees of freedom and to enable us to understand the complicated structure of tensor. Tucker decomposition can fulfill this requirement by expanding a matrix with a produce sum of a small tensor and matrices. It is very usual that small core tensor as well as small matrices can represent the original tensor well. Although there are multiple ways to compute Tucker decomposition, we employed Higher-order singular value decomposition (HOSVD) to perform Tucker decomposition. HOSVD is a simple algorithm since it performs singular value decomposition to a matrix generated from a tensor by unfolding. Thus Tucker decomposition with HOSVD is easy to perform and fitted to be applied to massive HOSVD data set.

### 5.4 TD-based unsupervised FE applied to the synthetic and Hi-C data

Higher-order singular value decomposition (HOSVD) [11] was applied to *x*_*ii*′*k*_ ∈ ℝ ^*N*×*N*×*K*^ to obtain

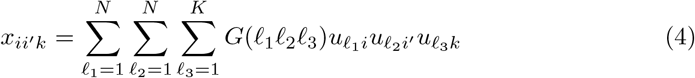

where *G* ∈ℝ ^*N ×N ×K*^ is a core tensor that represents the contribution of 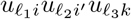 to 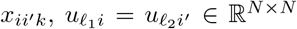 and 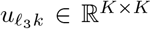 are singular value and orthogonal matrices. HOSVD was performed by applying `irlba` [26] to unfolded tensor *x*_*i*(*i*′*k*)_ ∈ℝ ^*N ×*(*NK*)^ in sparse matrix format using the Matrix package [27] since *x*_*ii*′*k*_ was too large to be stored in a dense matrix format. Because HOSVD is a set of singular value decompositions (SVD) applied to an unfolded tensor [11], HOSVD can be easily replaced with SVD. 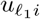 of interest was selected either based on its correlation with *y*_*i*_ (for the synthetic data) or with known functional sites (for the Hi-C dataset). Once 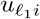 of interest was selected, *P* -values were attributed to *i* as

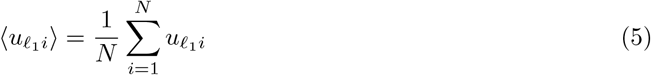

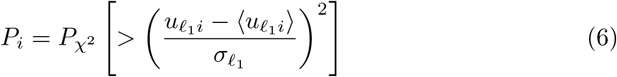

where 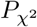 [*> x*] is the cumulative *χ* probability distribution, in which the argument is larger than *x* and 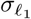 is the standard deviation of 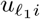,computed by optimization. The histogram of *P*_*i*_ was assumed to be flat if 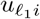 followed a Gaussian distribution. The optimization of 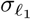 was performed such that the histogram of *P*_*i*_ was as flat as possible. To this end, first, the frequency of *P*_*i*_ in the *s*th bin, *h*_*s*_, 1 *≤s ≤S*, was computed by excluding *i*s whose associated adjusted *P* -values (see below) were less than the threshold value because *i*s whose associated adjusted *P* -values were less than the threshold value were expected to not obey a Gaussian distribution. Without excluding *i*s whose associated adjusted *P* -values were less than the threshold value, 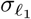 and *P*_*i*_ would be overestimated. Therefore, the number of *i*s whose associated adjusted *P* -values were less than the threshold value would be wrongly reduced, as well as those of the selected *i*s. Next, the standard deviation of *h*_*s*_, *σ*_*h*_, was computed as

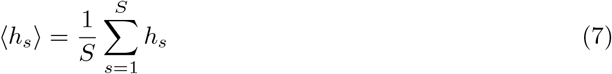

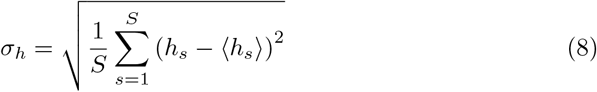

Then, 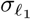 that minimized *σ*_*h*_ was computed, and *P*_*i*_ was computed using 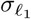, It is worth noting that this amounts to a self-consistency computation because adjusted *P* -values must be computed in order to exclude those *i*s associated with adjusted *P* -values below the threshold value to exclude these *i*s from the computation of *h*_*s*_. This process was iterated until the minimization of *σ*_*h*_ was complete. *P*_*i*_s were finally adjusted by the BH criterion [11] and those *i*s associated with adjusted *P* -values less than the threshold value were selected. In the case that the readers are interested in the processes included in TD-based unsupervised FE, please refer to our recently published textbook [11], or to the vignettes of two Bioconductor Packages [28, 29].

#### 5.4.1 Mean vs SD ratio

To quantify the differences of the selected *x*_*ii*′*k*_ between dataset (*k*) within a set of selected *i*s, Ω_*i*_, the Mean vs SD ratio was defined as

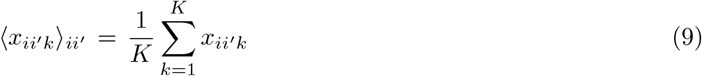

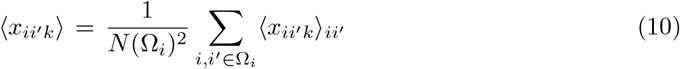

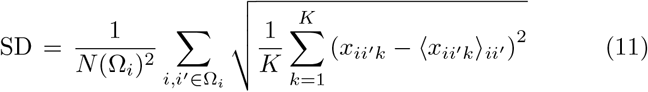

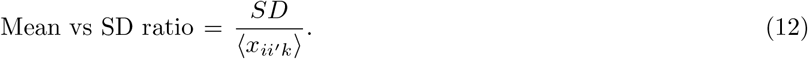

The summation of *i, i*^*′*^ was considered in Ω_*i*_ and *N* (Ω_*i*_) was used to denote the number of *i*s in Ω_*i*_.

### 5.3 Functional sites used to compute the correlation with 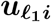 s or ⟨*x*_*ii*_*′k*⟩_*i*_ for the Hi-C datasets

#### 5.3.1 Overlap with bins

Overlaps between bins were computed using the `findOverlaps` function in IRanges [30].

#### 5.3.2 CTCF

The CTCF profile was retrieved from the AH104727 data in the AnnotationHub [31].

#### 5.3.3 PLS, pELS, and dELS

The PLS, pELS, and dELS profiles were obtained from 41586 2020 2493 MOESM12 ESM.txt [32].

### 5.4 Similarity

Similarity in Table 7 is defined as the Jaccard coefficient of tads domains.bed computed by hicFindTADs command in HiCExplorer between bin sizes 1,000,000 and 150,000; the Jaccard coefficient between distinct bin sizes is often used as the quality assessment of Hi-C profiles [33]. Computation was done by jaccard command in bedtools [34] as `bedtools jaccard -a file1 -b file2` where file1 and file2 are bed files for bin size 1,000,000 and 150,000, respectively.

## Declarations

## Ethics approval and consent to participate

Not applicable

## Data availability

We have gotten all the data analysed in this study from GSE260760 and GSE255264.

## Code availability

The code used to implement our method is publicly available from the GitHub repository at https://github.com/tagtag/TDbasedUFEHiC.

## Author contribution

Y. H. T. planned the study and performed the analyses. Y.-H.T. and T.T. evaluated the results and wrote and reviewed the manuscript. All the authors have read and agreed to the published version of the manuscript. The conceptualization, data curation, analysis were performed by Y. H. T.

## References

[1] Li, A., Yin, X., Xu, B., Wang, D., Han, J., Wei, Y., Deng, Y., Xiong, Y., Zhang, Z.: Decoding topologically associating domains with ultra-low resolution Hi-C data by graph structural entropy. Nature Communications 9(1), 3265 (2018) 10.1038/s41467-018-05691-7

[2] Sauria, M.E., Taylor, J.: Quasar: Quality assessment of spatial arrangement reproducibility in hi-c data. bioRxiv (2017) 10.1101/204438 https://www.biorxiv.org/content/early/2017/11/14/204438.full.pdf

[3] Zhang, Y., An, L., Xu, J., Zhang, B., W, J.Z., Hu, M., Tang, J., Yue, F.: Enhancing Hi-C data resolution with deep convolutional neural network HiCPlus. Nature Communications 9(1), 750 (2018) 10.1038/s41467-018-03113-2

[4] Yardimci, G.G., Ozadam, H., Sauria, M.E.G., Ursu, O., Yan, K.-K., Yang, T., Chakraborty, A., Kaul, A., Lajoie, B.R., Song, F., Zhan, Y., Ay, F., Gerstein, M., Kundaje, A., Li, Q., Taylor, J., Yue, F., Dekker, J., Noble, W.S.: Measuring the reproducibility and quality of Hi-C data. Genome Biology 20(1), 57 (2019) 10.1186/s13059-019-1658-7

[5] Yang, T., Zhang, F., Yardımcı, G.G., Song, F., Hardison, R.C., Noble, W.S., Yue, F., Li, Q.: Hicrep: assessing the reproducibility of hi-c data using a stratum-adjusted correlation coefficient. Genome Research 27(11), 1939–1949 (2017) 10.1101/gr.220640.117 http://genome.cshlp.org/content/27/11/1939.full.pdf+html

[6] Ursu, O., Boley, N., Taranova, M., Wang, Y.X.R., Yardimci, G.G., Stafford Noble, W., Kundaje, A.: GenomeDISCO: a concordance score for chromosome conformation capture experiments using random walks on contact map graphs. Bioinformatics 34(16), 2701–2707 (2018) 10.1093/bioinformatics/bty164 https://academic.oup.com/bioinformatics/article-pdf/34/16/2701/48917757/bioinformatics34162701.pdf

[7] Yan, K.-K., Yardımcı, G.G., Yan, C., Noble, W.S., Gerstein, M.: HiC-spector: a matrix library for spectral and reproducibility analysis of Hi-C contact maps. Bioinformatics 33(14), 2199–2201 (2017) 10.1093/bioinformatics/btx152 https://academic.oup.com/bioinformatics/article-pdf/33/14/2199/50314921/bioinformatics33142199.pdf

[8] Zou, C., Zhang, Y., Ouyang, Z.: HSA: integrating multi-track Hi-C data for genome-scale reconstruction of 3D chromatin structure. Genome Biology 17(1), 40 (2016) 10.1186/s13059-016-0896-1

[9] Stansfield, J.C., Cresswell, K.G., Dozmorov, M.G.: multiHiCcompare: joint normalization and comparative analysis of complex Hi-C experiments. Bioinformatics 35(17), 2916–2923 (2019) 10.1093/bioinformatics/btz048 https://academic.oup.com/bioinformatics/article-pdf/35/17/2916/50719885/bioinformatics35172916.pdf

[10] Li, A., Zeng, G., Wang, H., Li, X., Zhang, Z.: Dedoc2 identifies and characterizes the hierarchy and dynamics of chromatin tad-like domains in the single cells. Advanced Science 10(20), 2300366 (2023) 10.1002/advs.202300366 https://onlinelibrary.wiley.com/doi/10.1002/advs.202300366

[11] Taguchi, Y.-h.: Unsupervised Feature Extraction Applied to Bioinformatics: A PCA Based and TD Based Approach, 2nd edn. Unsupervised and Semi-Supervised Learning. Springer, Switzland (2024)

[12] Taguchi, Y.H., Turki, T.: Novel artificial intelligence-based identification of drug-gene-disease interaction using protein-protein interaction. BMC Bioinformatics 25(1), 377 (2024)

[13] Taguchi, Y.-h., Turki, T.: Advanced tensor decomposition-based integrated analysis of protein-protein interaction with cancer gene expression can improve coincidence with clinical labels. In: 2023 11th International Conference on Bioinformatics and Computational Biology (ICBCB), pp. 19–25 (2023). 10.1109/ICBCB57893.2023.10246633

[14] Taguchi, Y.-H., Turki, T.: Integrated analysis of gene expression and protein–protein interaction with tensor decomposition. Mathematics 11(17) (2023) 10.3390/math11173655

[15] Liu, T., Zhou, J., Chen, Y., Fang, J., Liu, S., Frangou, C., Wang, H., Zhang, J.: Genome-wide characterization of taz binding sites in mammary epithelial cells. Cancers 15(19) (2023) 10.3390/cancers15194713

[16] Li, Y., Liu, Y., Guo, Y.-Z., Liao, X.-F., Hu, B., Yu, T.: Spatio-temporal-spectral hierarchical graph convolutional network with semisupervised active learning for patient-specific seizure prediction. IEEE Transactions on Cybernetics 52(11), 12189–12204 (2022) 10.1109/TCYB.2021.3071860

[17] Cui, W., Xiang, Y., Wang, Y., Yu, T., Liao, X.-F., Hu, B., Li, Y.: Deep multiview module adaption transfer network for subject-specific eeg recognition. IEEE Transactions on Neural Networks and Learning Systems, 1–14 (2024) 10.1109/TNNLS.2024.3350085

[18] Shi, B., Chen, J., Chen, H., Lin, W., Yang, J., Chen, Y., Wu, C., Huang, Z.: Prediction of recurrent spontaneous abortion using evolutionary machine learning with joint self-adaptive sime mould algorithm. Computers in Biology and Medicine 148, 105885 (2022) 10.1016/j.compbiomed.2022.105885

[19] Lu, W., Zhao, H., He, Q., Huang, H., Jin, X.: Category-consistent deep network learning for accurate vehicle logo recognition. Neurocomputing 463, 623–636 (2021) 10.1016/j.neucom.2021.08.030

[20] Wolff, J., Rabbani, L., Gilsbach, R., Richard, G., Manke, T., Backofen, R., Grüning, B.A.: Galaxy hicexplorer 3: a web server for reproducible hi-c, capture hi-c and single-cell hi-c data analysis, quality control and visualization. Nucleic Acids Research 48(W1), 177–184 (2020) 10.1093/nar/gkaa220 https://academic.oup.com/nar/article-pdf/48/W1/W177/33433206/gkaa220.pdf

[21] Bittner, N., Shi, C., Zhao, D., Ding, J., Southam, L., Swift, D., Kreitmaier, P., Tutino, M., Stergiou, O., Cheung, J.T.S., Katsoula, G., Hankinson, J., Wilkinson, J.M., Orozco, G., Zeggini, E.: Primary osteoarthritis chondrocyte map of chromatin conformation reveals novel candidate effector genes. Annals of the Rheumatic Diseases 83(8), 1048–1059 (2024) 10.1136/ard-2023-224945 https://ard.bmj.com/content/83/8/1048.full.pdf

[22] Nolan, B., Harris, H.L., Kalluchi, A., Reznicek, T.E., Cummings, C.T., Rowley, M.J.: Hicrayon reveals distinct layers of multi-state 3d chromatin organization. bioRxiv (2024) 10.1101/2024.02.11.579821 https://www.biorxiv.org/content/early/2024/02/12/2024.02.11.579821.full.pdf

[23] Harris, H.L., Gu, H., Olshansky, M., Wang, A., Farabella, I., Eliaz, Y., Kalluchi, A., Krishna, A., Jacobs, M., Cauer, G., Pham, M., Rao, S.S.P., Dudchenko, O., Omer, A., Mohajeri, K., Kim, S., Nichols, M.H., Davis, E.S., Gkountaroulis, D., Udupa, D., Aiden, A.P., Corces, V.G., Phanstiel, D.H., Noble, W.S., Nir, G., Pierro, M.D., Seo, J.-S., Talkowski, M.E., Aiden, E.L., M, J.R.: Chromatin alternates between A and B compartments at kilobase scale for subgenic organization. Nature Communications 14(1), 3303 (2023) 10.1038/s41467-023-38429-1

[24] Leinonen, R., Sugawara, H., Shumway, o.b.o.t.I.N.S.D.C. Martin: The Sequence Read Archive. Nucleic Acids Research 39(suppl 1), 19–21 (2010) 10.1093/nar/gkq1019 https://academic.oup.com/nar/article-pdf/39/suppl1/D19/7624335/gkq1019.pdf

[25] Servant, N., Varoquaux, N., Lajoie, B.R., Viara, E., Chen, C.-J., Vert, J.-P., Heard, E., Dekker, J., Barillot, E.: HiC-Pro: an optimized and flexible pipeline for Hi-C data processing. Genome Biology 16(1), 259 (2015) 10.1186/s13059-015-0831-x

[26] Baglama, J., Reichel, L., Lewis, B.W.: Irlba: Fast Truncated Singular Value Decomposition and Principal Components Analysis for Large Dense and Sparse Matrices. (2022). R package version 2.3.5.1. https://CRAN.R-project.org/package=irlba

[27] Bates, D., Maechler, M., Jagan, M.: Matrix: Sparse and Dense Matrix Classes and Methods. (2024). R package version 1.7-0. https://CRAN.R-project.org/package=Matrix

[28] Taguchi, Y.-h.: TDbasedUFE: Tensor Decomposition Based Unsupervised Feature Extraction. (2023). 10.18129/B9.bioc.TDbasedUFE. R package version 1.0.0. https://bioconductor.org/packages/TDbasedUFE

[29] Taguchi, Y.-h.: TDbasedUFEadv Advanced Package of Tensor Decomposition Based Unsupervised Feature Extraction. (2023). 10.18129/B9.bioc.TDbasedUFEadv. R package version 1.0.0. https://bioconductor.org/packages/TDbasedUFEadv

[30] Lawrence, M., Huber, W., Pagès, H., Aboyoun, P., Carlson, M., Gentleman, R., Morgan, M., Carey, V.: Software for computing and annotating genomic ranges. PLoS Computational Biology 9 (2013) 10.1371/journal.pcbi.1003118

[31] Morgan, M., Shepherd, L.: AnnotationHub: Client to Access AnnotationHub Resources. (2023). 10.18129/B9.bioc.AnnotationHub. R package version 3.8.0. https://bioconductor.org/packages/AnnotationHub

[32] Abascal, F., Acosta, R., Addleman, N.J., Adrian, J., Afzal, V., Ai, R., Aken, B., Akiyama, J.A., Jammal, O.A., Amrhein, H., Anderson, S.M., Andrews, G.R., Antoshechkin, I., Ardlie, K.G., Armstrong, J., Astley, M., Banerjee, B., Barkal, A.A., Barnes, I.H.A., Barozzi, I., Barrell, D., Barson, G., Bates, D., Baymuradov, U.K., Bazile, C., Beer, M.A., Beik, S., M, A.B., Bennett, R., Bouvrette, L.P.B., Bernstein, B.E., Berry, A., Bhaskar, A., Bignell, A., Blue, S.M., Bodine, D.M., Boix, C., Boley, N., Borrman, T., Borsari, B., Boyle, A.P., Brandsmeier, L.A., Breschi, A., Bresnick, E.H., Brooks, J.A., Buckley, M., Burge, C.B., Byron, R., Cahill, E., Cai, L., Cao, L., Carty, M., Castanon, R.G., Castillo, A., Chaib, H., Chan, E.T., Chee, D.R., Chee, S., Chen, H., Chen, H., Chen, J.-Y., Chen, S. J M.C., Chhetri, S.B., Choudhary, J.S., Chrast, J., Chung, D., Clarke, D., Cody, N.A.L., Coppola, C.J., Coursen, J., D’Ippolito, A.M., Dalton, S., Danyko, C., Davidson, C., Davila-Velderrain, J., Davis, C.A., Dekker, J., Deran, A., DeSalvo, G., Despacio-Reyes, G., Dewey, C.N., Dickel, D.E., Diegel, M., Diekhans, M., Dileep, V., Ding, B., Djebali, S., Dobin, A., Dominguez, D., Donaldson, S., Drenkow, J., Dreszer, T.R., Drier, Y., Duff, M.O., Dunn, D., Eastman, C., Ecker, J.R., Edwards, M.D., El-Ali, N., Elhajjajy, S.I., Elkins, K., Emili, A., Epstein, C.B., Evans, R.C., Ezkurdia, I., Fan, K., Farnham, P.J., Farrell, N.P., Feingold, E.A., Ferreira, A.-M., Fisher-Aylor, K., Fitzgerald, S., Flicek, P., Foo, C.S., Fortier, K., Frankish, A., Freese, P., Fu, S., Fu, X.-D., Fu, Y., Fukuda-Yuzawa, Y., Fulciniti, M., Funnell, A.P.W., Gabdank, I., Galeev, T., Gao, M., Giron, C.G., Garvin, T.H., Gelboin-Burkhart, C.A., Georgolopoulos, G., Gerstein, M.B., Giardine, B.M., Gifford, D.K., Gilbert, D.M., Gilchrist, D.A., Gillespie, S., Gingeras, T.R., Gong, P., Gonzalez, A., Gonzalez, J.M., Good, P., Goren, A., Gorkin, D.U., Graveley, B.R., Gray, M., Greenblatt, J.F., Griffiths, E., Groudine, M.T., Grubert, F., Gu, M., Guigó, R., Guo, H., Guo, Y., Zheng, Y., Gursoy, G., Gutierrez-Arcelus, M., Halow, J., Hardison, R.C., Hardy, M., Hariharan, M., Harmanci, A., Harrington, A., Harrow, J.L., Hashimoto, T.B., Hasz, R.D., Hatan, M., Haugen, E., Hayes, J.E., He, P., He, Y., Heidari, N., Hendrickson, D., Heuston, E.F., Hilton, J.A., Hitz, B.C., Hochman, A., Holgren, C., Hou, L., Hou, S., Hsiao, Y.-H.E., Hsu, S., Huang, H., Hubbard, T.J., Huey, J., Hughes, T.R., Hunt, T., Ibarrientos, S., Issner, R., Iwata, M., Izuogu, O., Jaakkola, T., Jameel, N., Jansen, C., Jiang, L., Jiang, P., Johnson, A., Johnson, R., Jungreis, I., Kadaba, M., Kasowski, M., Kasparian, M., Kato, M., Kaul, R., Kawli, T., Kay, M., Keen, J.C., Keles, S., Keller, C.A., Kelley, D., Kellis, M., Kheradpour, P., Kim, D.S., Kirilusha, A., Klein, R.J., Knoechel, B., Kuan, S., Kulik, M.J., Kumar, S., Kundaje, A., Kutyavin, T., Lagarde, J., Lajoie, B.R., Lambert, N.J., Lazar, J., Lee, A.Y., Lee, D., Lee, E., Lee, J.W., Lee, K., Leslie, C.S., Levy, S., Li, B., Li, H., Li, N., Li, S., Li, X., Li, Y., Li, Y., Li, Y., Li, Y., Lian, J., Libbrecht, M.W., Lin, S., Lin, Y., Liu, D., Liu, J., Lu, A., Liu, T. X S.L., Liu, Y., Liu, Y., Long, M., Lou, S., Loveland, J., Lu, A., Lu, Y., Lècuyer, E., Ma, L., Mackiewicz, M., Mannion, B.J., Mannstadt, M., Manthravadi, D., Marinov, G.K., Martin, F.J., Mattei, E., McCue, K., McEown, M., McVicker, G., Meadows, S.K., Meissner, A., Mendenhall, E.M., Messer, C.L., Meuleman, W., Meyer, C., Miller, S., Milton, M.G., Mishra, T., Moore, D.E., Moore, H.M., Moore, J.E., Moore, S.H., Moran, J., Mortazavi, A., Mudge, J.M., Munshi, N., Murad, R., Myers, R.M., Nandakumar, V., Nandi, P., Narasimha, A.M., Narayanan, A.K., Naughton, H., Navarro, F.C.P., Navas, P., Nazarovs, J., Nelson, J., Neph, S., Neri, F.J., Nery, J.R., Nesmith, A.R. J S.N., Newberry, K.M., Ngo, V., Nguyen, R., Nguyen, T.B., Nguyen, T., Nishida, A., Noble, W.S., Novak, C.S., Novoa, E.M., Nuñez, B., O’Donnell, C.W., Olson, S., Onate, K.C., Otterman, E., Ozadam, H., Pagan, M., Palden, T., Pan, X., Park, Y., E, C.P., Paten, B., Pauli-Behn, F., Pazin, M.J., Pei, B., Pennacchio, L.A., Perez, A.R., Perry, E.H., Pervouchine, D.D., Phalke, N.N., Pham, Q., Phanstiel, D.H., Plajzer-Frick, I., Pratt, G.A., Pratt, H.E., Preissl, S., Pritchard, J.K., Pritykin, Y., Purcaro, M.J., Qin, Q., Quinones-Valdez, G., Rabano, I., Radovani, E., Raj, A., Rajagopal, N., Ram, O., Ramirez, L., Ramirez, R.N., Rausch, D., Raychaudhuri, S., Raymond, J., Razavi, R., Reddy, T.E., Reimonn, T.M., Ren, B., Reymond, A., Reynolds, A., Rhie, S.K., Rinn, J., Rivera, M., Rivera-Mulia, J.C., Roberts, B.S., Rodriguez, J.M., Rozowsky, J., Ryan, R., Rynes, E., Salins, D.N., Sandstrom, R., Sasaki, T., Sathe, S., Savic, D., Scavelli, A., Scheiman, J., Schlaffner, C., Schloss, J.A., Schmitges, F.W., See, L.H., Sethi, A., Setty, M., Shafer, A., Shan, S., Sharon, E., Shen, Q., Shen, Y., Sherwood, R.I., Shi, M., Shin, S., Shoresh, N., Siebenthall, K., Sisu, C., Slifer, T., Sloan, C.A., Smith, A., Snetkova, V., Snyder, M.P., Spacek, D.V., Srinivasan, S., Srivas, R., Stamatoyannopoulos, G., Stamatoyannopoulos, J.A., Stanton, R., Steffan, D., Stehling-Sun, S., J, S.S., Su, A., Sundararaman, B., Suner, M.-M., Syed, T., Szynkarek, M., Tanaka, F.Y., Tenen, D., Teng, M., Thomas, J.A., Toffey, D., Tress, M.L., Trout, D.E., Trynka, G., Tsuji, J., Upchurch, S.A., Ursu, O., Uszczynska-Ratajczak, B., Uziel, M.C., Valencia, A., Biber, B.V., Velde, A.G.v.d., Nostrand, E.L.V., Vaydylevich, Y., Vazquez, J., Victorsen, A., Vielmetter, J., Vierstra, J., Visel, A., Vlasova, A., Vockley, C.M., Volpi, S., Vong, S., Wang, H., Wang, M., Wang, Q., Wang, R., Wang, T., Wang, W., Wang, X., Wang, Y., Watson, N.K., Wei, X., Wei, Z., Weisser, H., Weissman, S.M., Welch, R., Welikson, R.E., Weng, Z., Westra, H.-J., Whitaker, J.W., White, C., White, K.P., Wildberg, A., Williams, B.A., Wine, D., Witt, H.N., Wold, B., Wolf, M., Wright, J., Xiao, R., Xiao, X., Xu, J., Xu, J., Yan, K.-K., Yan, Y., Yang, H., Yang, X., Yang, Y.-W., Yardımcı, G.G., Yee, B.A., Yeo, G.W., Young, T., Yu, T., Yue, F., Zaleski, C., Zang, C., Zeng, H., Zeng, W., Zerbino, D.R., Zhai, J., Zhan, L., Zhan, Y., Zhang, B., Zhang, J., Zhang, J., Zhang, K., Zhang, L., Zhang, P., Zhang, Q., Zhang, X.-O., Zhang, Y., Zhang, Z., Zhao, Y., Zheng, Y., Zhong, G., Zhou, X.-Q., Zhu, Y., Zimmerman, J., Moore, J.E., Purcaro, M.J., Pratt, H.E., Epstein, C.B., Shoresh, N., Adrian, J., Kawli, T., Davis, C.A., Dobin, A., Kaul, R., Halow, J., Nostrand, E.L.V., Freese, P., Gorkin, D.U., Shen, Y., He, Y., Mackiewicz, M., Pauli-Behn, F., Williams, B.A., Mortazavi, A., Keller, C.A., Zhang, X.-O., Elhajjajy, S.I., Huey, J., Dickel, D.E., Snetkova, V., Wei, X., Wang, X., Rivera-Mulia, J.C., Rozowsky, J., Zhang, J., Chhetri, S.B., Zhang, J., Victorsen, A., White, K.P., Visel, A., Yeo, G.W., Burge, C.B., Lècuyer, E., Gilbert, D.M., Dekker, J., Rinn, J., Mendenhall, E.M., Ecker, J.R., Kellis, M., Klein, R.J., Noble, W.S., Kundaje, A., Guigó, R., Farnham, P.J., J, M.C., Myers, R.M., Ren, B., Graveley, B.R., Gerstein, M.B., Pennacchio, L.A., Snyder, M.P., Bernstein, B.E., Wold, B., Hardison, R.C., Gingeras, T.R., Stamatoyannopoulos, J.A., Weng, Z.: Expanded encyclopaedias of DNA elements in the human and mouse genomes. Nature 583(7818), 699–710 (2020) 10.1038/s41586-020-2493-4

[33] Forcato, M., Nicoletti, C., Pal, K., Livi, C.M., Ferrari, F., Bicciato, S.: Comparison of computational methods for hi-c data analysis. Nature Methods 14(7), 679–685 (2017) 10.1038/nmeth.4325

[34] Quinlan, A.R., Hall, I.M.: Bedtools: a flexible suite of utilities for comparing genomic features. Bioinformatics 26(6), 841–842 (2010) 10.1093/bioinformatics/btq033

